# Crossing fitness valleys via double substitutions within codons

**DOI:** 10.1101/511469

**Authors:** Frida Belinky, Itamar Sela, Igor B. Rogozin, Eugene V. Koonin

## Abstract

Single nucleotide substitutions in protein-coding genes can be divided into synonymous (S), with little fitness effect, and non-synonymous (N) ones that alter amino acids and thus generally have a greater effect. Most of the N substitutions are affected by purifying selection that eliminates them from evolving populations. However, additional mutations of nearby bases can modulate the deleterious effect of single substitutions and thus might be subject to positive selection. To elucidate the effects of selection on double substitutions in all codons, it is critical to differentiate selection from mutational biases. We approached this problem by comparing the fractions of double substitutions within codons to those of the equivalent double S substitutions in adjacent codons. Under the assumption that substitutions occur one at a time, all within-codon double substitutions can be represented as “ancestral-intermediate-final” sequences and can be partitioned into 4 classes: 1) SS: S intermediate – S final, 2) SN: S intermediate – N final, 3) NS: N intermediate – S final, 4) NN: N intermediate – N final. We found that the selective pressure on the second substitution markedly differs among these classes of double substitutions. Analogous to single S substitutions, SS evolve neutrally whereas, analogous to single N substitutions, SN are subject to purifying selection. In contrast, NS show positive selection on the second step because the original amino acid is recovered. The NN double substitutions are heterogeneous and can be subject to either purifying or positive selection, or evolve neutrally, depending on the amino acid similarity between the final or intermediate and the ancestral states. The general trend is that the second mutation compensates for the deleterious effect of the first one, resulting in frequent crossing of valleys on the fitness landscape.

## Introduction

In classic population genetics, mutations are assumed to occur one at a time, independently of each other^1-5^. However, clustering of mutations, in particular, those occurring in adjacent sites (multi-nucleotide mutations) has been documented in many diverse organisms^6-13^. Multi-nucleotide substitutions potentially could originate from mutational biases, selection, or a combination of both. Recently, it has been claimed that positive selection is over-estimated by the branch-site test (BST) because many if not most of the sites supporting positive selection actually are multi-nucleotide substitutions that could result from multi-nucleotide mutations^14^. However, independent of BST, double substitutions within the same codon in protein-coding genes have been repeatedly claimed to be driven by positive selection. This conclusion follows from the comparison of the observed frequencies of double substitutions to those expected from the frequencies of single substitutions. If the frequency of a double substitution is significantly greater than the product of the frequencies of the respective single substitutions, positive selection is inferred^15-17^. Such apparent signs of positive selection affecting double substitutions have been detected as a general trend in the mouse-rat lineage^15^. Similar conclusions have been reached for double substitutions in codons for serine, the only amino acid that is encoded by two disjoint series of codons. In the case of serine, the proposed scenario is that a non-synonymous (N) substitution that leads to the replacement of a serine with another amino acid and is hence deleterious is followed by a second substitution that restores serine and, accordingly, the protein function and the original fitness value^17^. The fixation of the second mutation has been attributed to positive selection, and the observed excessive frequency of double substitutions has been explained by this effect of selection, as opposed to a mutational bias.

Similarly, signatures of positive selection have been found for double substitutions in stop codons in bacteria (UAG>UGA and UGA>UAG), which could be attributed to the deleterious, non-stop intermediate state, UGG^16^. Furthermore, slightly advantageous back mutations are expected under the nearly neutral model^18^. Thus, a second mutation in a codon that reverts a non-synonymous substitution to restore the codon for the original amino acid, generally, is expected to be advantageous. However, given that the apparent positive selection in codon double substitutions could be potentially explained by biased mutational processes that favor multi-nucleotide substitutions^6,9,14,17,19^, it is essential to compare codon double substitutions to an appropriate null model in order to accurately infer selection.

Following the well-established principles of identification of selective pressure by comparison of non-synonymous to synonymous rates^20-26^, to assess the selection that affects double substitutions within codons, we compared the double fraction (DF) of each such double substitution to the DF of adjacent equivalent double synonymous substitutions. We categorize codon double substitution into four classes and show that these classes of codon double substitution are associated with different types of selection acting on the second substitution step.

## Results

### Inference of selection on codon double substitutions by comparison to null models

From triplets of genomes with reliable phylogenetic relationships that were extracted from the ATGC database^17,27^, we obtained frequencies of double and single substitutions in codons, and in double synonymous controls (see Methods for details). The key difference between the present work and the previous studies is that all the analyses included comparison to double synonymous substitutions that served as null models for the double substitutions in codons. Although it is well known that transition and transversion rates differ substantially^22,28,29^, it is unclear to what extent the adjacency of mutations is affected by base composition. For example, DNA polymerase ƞ tends to produce an excessive amount of simultaneous double transitions in A/T-rich context^30^ whereas DNA polymerase ζ frequently produces transversions in C/G-rich context^31,32^. Another important issue is the balance between consecutive double substitutions (independent stepwise fixation of adjacent mutations) and simultaneous double substitutions. This issue cannot be ignored because some replication enzymes are known to produce or initiate production of excessive amounts of simultaneous double substitutions under certain conditions^33-37^. Therefore, we compared the frequencies of all codon double substitutions to all possible types of double synonymous substitutions that were captured in two null models (Fig. 1). The first null model (syn_31) included a synonymous substitution in the 3^rd^ position of a codon followed by another synonymous substitution in the 1^st^ position of the next codon. The second null model (syn_33) included non-adjacent synonymous substitutions in 3^rd^ codon positions of consecutive codons. We found that the double fraction (DF), i.e. the observed double substitution frequency divided by sum of the cumulative single substitution frequency and the double frequency (see Methods for further details) was typically higher for the syn_31 model compared to the syn_33 model suggesting the existence of a mutational bias in adjacent positions (Fig. 1). This difference was only statistically significant under the t-test but not with the non-parametric U-test.

**Figure 1.**
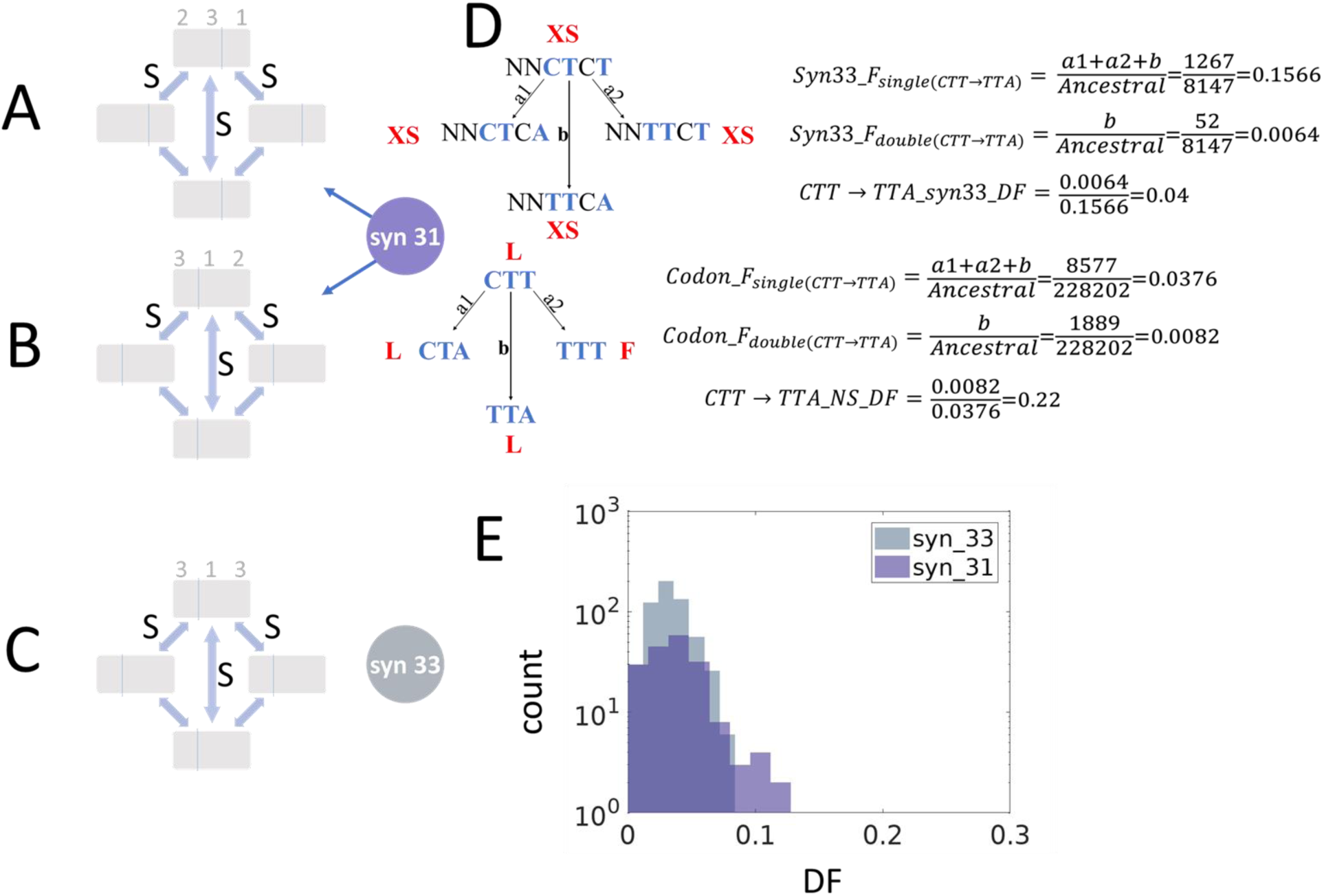
Double synonymous substitutions in adjacent codons used as null models and calculation of DF. (A) A constant 2nd codon positions followed by a 4-fold degenerate site in the 3rd codon positions which is followed by a 2-fold degenerate site in the 1st codon position of the next codon (B) A 4-fold degenerate site in the 3rd codon position followed by a 2-fold degenerate site in the 1st codon position of the next codon, which is followed by a constant base in the 2nd codon position of the second codon. (C) A 4-fold degenerate site in the 3rd codon position followed by a constant 1st codon position in the second codon of which the 2nd position is disregarded and by a 4-fold degenerate site in the 3rd codon position. (E) An example calculation of the DF under the null model syn_33 and an example calculation of the DF in an NS codon double substitution. (E) Comparison of DF between the two null models, syn_31 (adjacent synonymous substitutions) and syn_33 (non-adjacent synonymous substitutions). The difference between the two distributions is significant according to t-test (p-val=0.0038) but not significant with a Utest (p-val=0.104).

The DF is assumed to be proportional to the second step substitution rate. If the elevated DF of codon double substitutions results solely from a multi-nucleotide mutational bias, the comparison to the null model is expected to show no significant difference. Conversely, a significantly lower DF compared to that of the null model is indicative of purifying selection, whereas a significantly higher DF points to positive selection.

### Distinct selection regimes for different types of codon double substitutions

Representing all within-codon double substitutions in the general form, “ancestral-intermediate-final”, we define the following 4 combinations (Fig. 2A): 1) SS: S intermediate – S final, 2) SN: S intermediate – N final, 3) NS: N intermediate – S final, 4) NN: N intermediate – N final.

**Figure 2.**
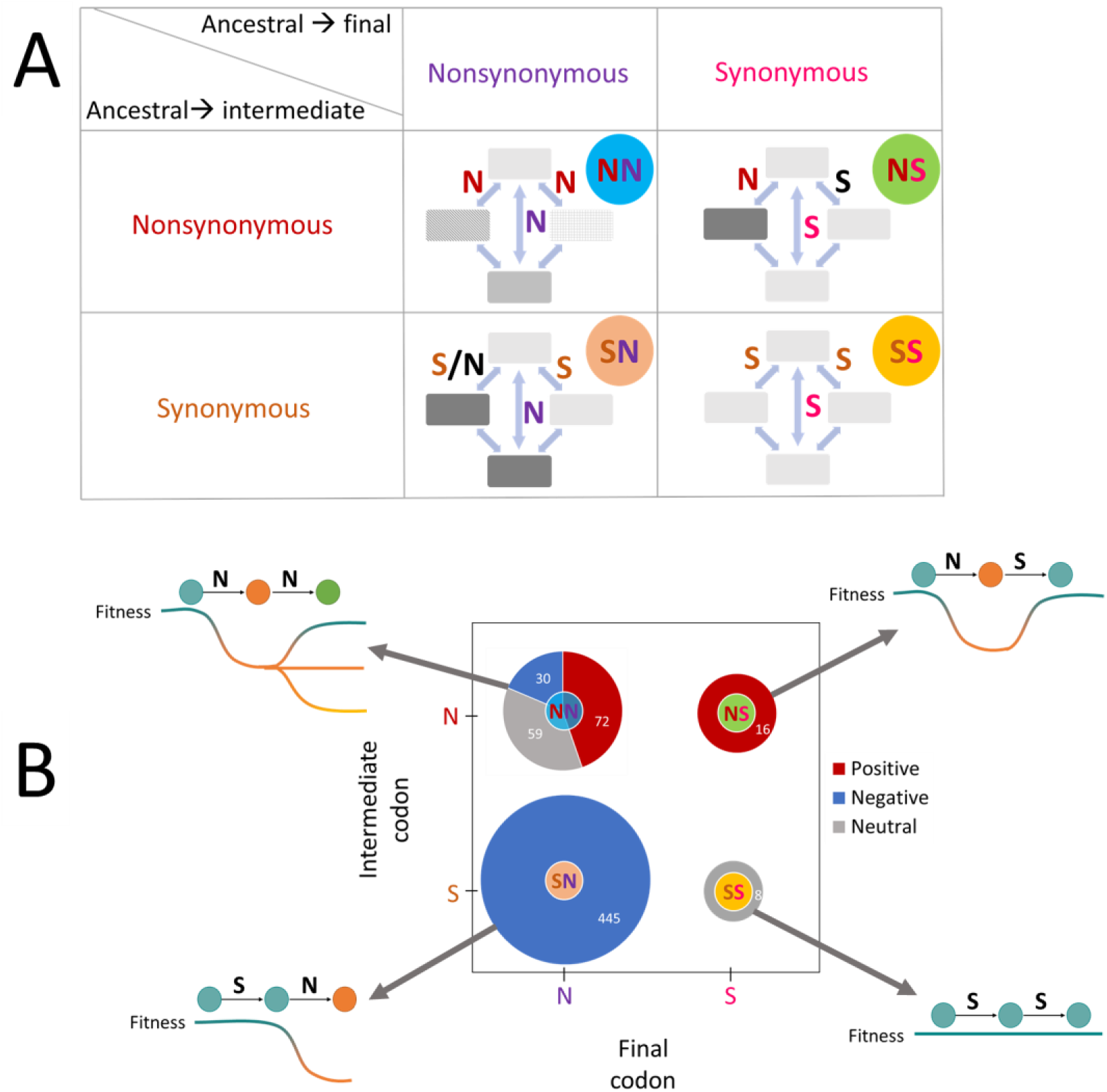
Classification of the double codon substitutions. (A) Four combinations of codon double substitutions based on synonymy of ancestral and derived (final) codons, and synonymy of intermediate state codons to the ancestral codons. (B) Selective pressure in different codon double substitutions classes. **Positive**, cases compatible with positive selection, where a codon double substitution has a significantly higher DF than the corresponding double synonymous substitution. **Negative**, cases compatible with purifying selection, where a codon double substitution has a significantly lower DF than the corresponding double synonymous substitution. **Neutral**, cases where the codon DF was not significantly different from that of the corresponding synonymous DF.

Additionally, we compare the DF between fast and slow evolving genes (see Methods). Changes that are subject to purifying selection are compatible with a higher DF in fast vs. slow evolving genes, and conversely, changes drive by positive selection are compatible with a higher DF in slow vs. fast evolving genes.

The results of these analyses reveal distinct selection regimes for the 4 classes of codon double substitutions (Fig. 2B). For the SS changes, neutrality cannot be rejected, the SN changes are subject to purifying selection, the NS changes are driven by positive selection, and NN changes exhibit a mixture of all three regimes depending on the similarity of the amino acids encoded by the intermediate and final codons to the original amino acid.

#### SS: double synonymous substitutions

For double synonymous substitutions, neutrality cannot be rejected by comparison to both null models using the U-test, which is the appropriate test for this small sample size (Fig. 3A). Thus, the DF values of the SS substitutions can be explained by the frequency of multi-nucleotide mutations suggesting that SS double substitutions evolve (nearly) neutrally, similar to single synonymous substitutions (Fig. S1). Additionally, there was no significant difference between the DF values of SS double substitutions in fast and slow evolving genes (Fig. 3A) which is compatible with the neutral evolutionary regime. Nevertheless, although the bulk analysis of the SS substitutions yields results compatible with neutrality, most of the individual SS cases seem to involve weak positive selection after the BH correction, which can be linked to codon bias (Fig. S2).

**Figure 3.**
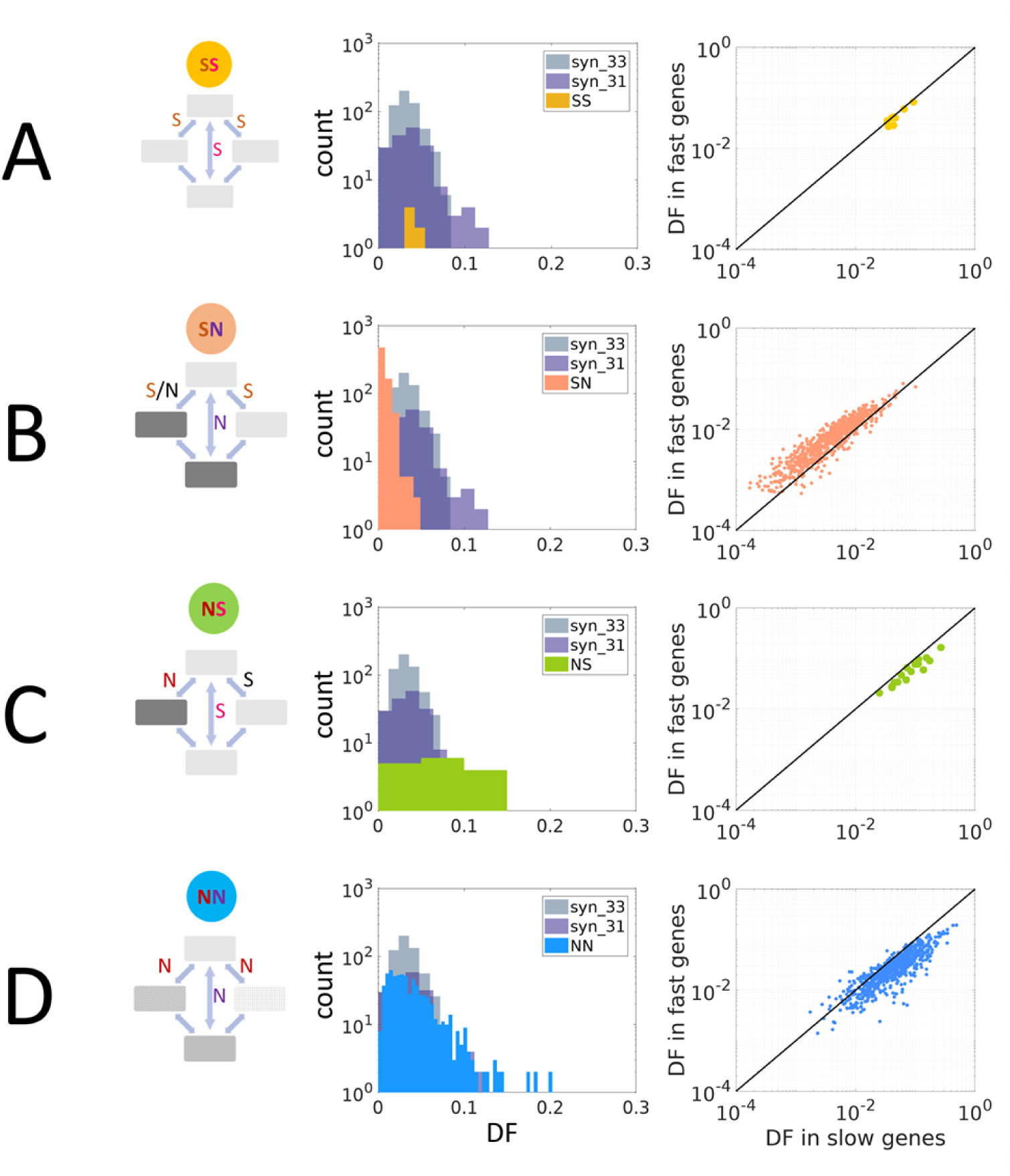
Selective regimes of the codon double substitutions. The panels on the left show the comparison of each codon double substitution class to the double synonymous null models, and the panels to the right show the comparisons between the DF of each of the classes in fast vs. slow evolving genes. (A) SS, double synonymous codon substitutions (B) SN, at least one synonymous intermediate codon, non-synonymous final codon (C) NS, one non-synonymous intermediate, synonymous final codon (D) NN – both intermediates and the final codon are non-synonymous to the ancestral.

#### SN: synonymous substitution followed by a non-synonymous one

The DF values for SN double substitutions are significantly lower than those for both syn_31 and syn_33 null models (Fig. 3B), indicating that the second step of these double substitutions is subject to purifying selection. Similarly to single non-synonymous substitutions (Fig. S1), the SN doubles show significantly higher DF values in fast evolving genes compared to slow evolving genes (Fig. 3B), which is also indicative of purifying selection. Analysis of individual cases of SN substitutions, after the BH correction, showed that 88% were compatible with purifying selection, for 10% neutrality could not be rejected, and less than 2% were compatible with positive selection (see Supplementary file for details).

#### NS: non-synonymous substitution followed by a synonymous one

The NS double substitutions show significantly higher DF values compared to both null models (Fig. 3C). This pattern is compatible with positive selection driving the second substitution which returns to the original amino acid state. The NS double substitutions also show higher DF in slow compared to fast evolving genes, which is compatible with positive selection (Fig. 3C). Analysis of individual NS double substitutions, after BH correction, resulted in 15 of the 16 cases that exhibit positive selection (93%). The only exception is TTG>CTC, for which the DF of the NS change was greater than that of the null model, but the difference was not statistically significant.

#### NN: double non-synonymous substitutions

For the NN double substitutions, detailed comparison of individual cases reveals a mixture of positive selection, purifying selection and neutral evolution. a. Neutrality cannot be rejected by the comparison of the DF values of NN doubles to the syn_31 null model. In contrast, the comparison between slow and fast evolving genes shows that DF of NN doubles is higher in slow compared to fast evolving genes, which is compatible with positive selection. Given this discrepancy between the results of the two tests, we performed an individual comparison for each NN change, with the same mutation types in the corresponding null model. This analysis of individual NN double substitutions (Table S1), after BH correction for multiple testing, demonstrated positive selection for 44% of the NN doubles, purifying selection for 19%, and neutral evolution for 36% (Fig. 3D).

### Modes of selection reflect amino-acid similarity

We hypothesized that the split of the NN into those evolving under positive selection, purifying selection or neutrally had to do with the (dis)similarity between the original, intermediate and final amino acid residues (Figure 1A). To test this hypothesis, we compared the differences in amino-acid similarity (DAS) between the subsets of NN, SN and NS doubles for which positive selection, purifying selection, or a neutral evolution regime were detected (Figure 4). This difference was calculated as DAS=S_of_-S_oi_ where S_of_ and S_oi_ are the similarity measures between the original and the final or intermediate amino acids, respectively. The measures of similarity between amino acid residues were extracted from 94 amino-acid similarity matrices that are available in the AAindex database^38^. For 85 of the 94 matrices, there was a significant difference between the DAS values of NN cases under positive selection compared to those under purifying selection. For most of the cases in the positively selected subset, DAS >0, i.e. the final amino acid is significantly more similar to the original than the intermediate amino acid. Conversely, for most of the cases in the negatively selected subset of the NN doubles, DAS <0, i.e. the second mutation decreases the similarity of the amino acid in the given position to the original one. Significant differences between positive vs. neutral, and neutral vs. negative subsets were observed as well albeit with fewer matrices (74 and 62, respectively). Focusing on 5 similarity/distance matrices that are based solely on psychochemical properties and thus rule out potential circular reasoning, we observed a significant difference between the DAS values for the NN cases under positive and purifying selection, and between the cases under positive selection and neutral evolution. However, the difference between the neutral cases and those under purifying selection was not significant.

**Figure 4.**
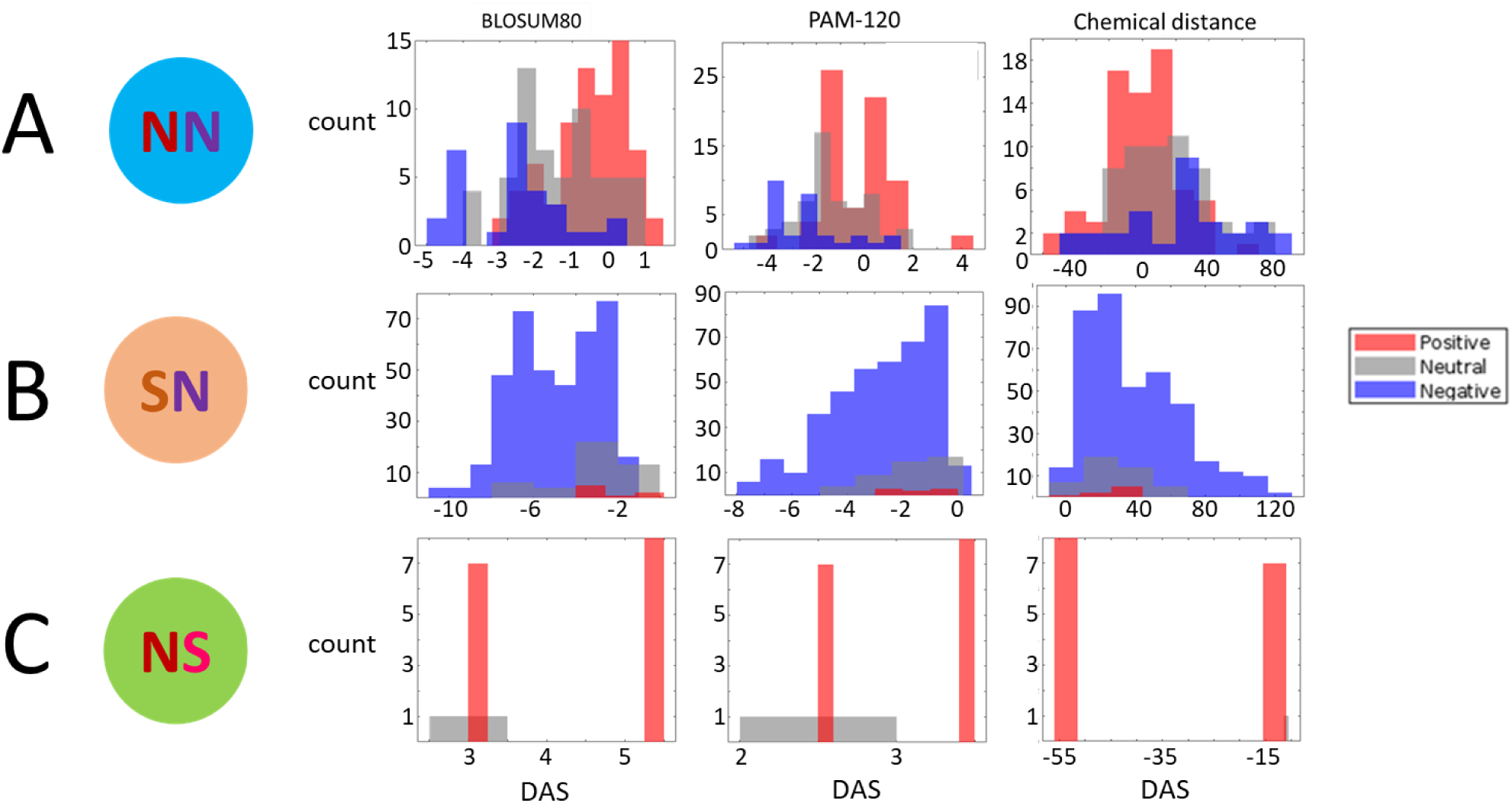
Similarity between the ancestral, intermediate and final amino acids for different classes of double substitutions. The DAS metric measures the difference in amino acid similarity/distance for the original→ final vs. original→ intermediate codons. DAS = AA similarity (original→final) – average AA similarity (original→intermediate). Three comparisons, using different amino acid similarity/distance matrices, are shown. (A) NN double substitutions (B) SN double substitutions (C) NS double substitutions.

We performed analogous comparisons also for the SN and NS classes of doubles substitutions. Although in each of the SN and NS cases, there is only one non-synonymous, with only two amino acids involved, the DAS values can be formally calculated by including the ancestral vs intermediate and the ancestral-final amino acid self-comparisons for SN and NS, respectively. For the SN cases, 78 of the 94 matrices yielded a significant difference between the negative and neutral groups, i.e. the final amino acid is less similar to the original one in the cases of purifying selection compared to neutral cases. For 56 matrices, there was a significant difference between the positive and negative groups, but only 5 matrices showed a significant difference between the positive and neutral groups. For the 5 similarity/distance matrices that are based solely on psychochemical properties, only the difference between the neutral and negative groups was significant. For the NS cases, there was no significant difference between the 15 positive cases and the single neutral case. The lack of statistical support in the latter comparisons is most likely due to the small number of positive cases for the SN class and, conversely, the dominance of positive selection in the NS class.

#### Additional controls for mutational biases

For the SN, NS, and NN doubles, similar results were obtained when transitions and transversions were analyzed separately (Fig. S3) and when double substitutions in non-coding regions were used as null-models instead of the syn_31 or syn_33 models (Fig. S4). The only exception were the SS doubles which had a greater DF compared to non-coding double substitutions (Fig. S3). This finding is likely explained by the purifying selection that, on average, affects non-coding regions to a greater extent than synonymous codon positions^39^.

#### Contribution of simultaneous double mutations

We estimated the frequency of simultaneous double mutations by calculating the difference between the observed double frequency in the null models, and the product of single synonymous substitutions (see methods). The estimated frequencies of the double mutations range from zero to 0.11, with the means of 0.0015 and 0.02 for syn33 and syn31, respectively. These frequencies are not negligible as they account for a mean of 53% of the double substitution frequency in syn33 and for 66% in syn31 (Fig. S5). Although the DF is not significantly different between syn33 and syn31, the estimated proportion of simultaneous mutations is (Fig. S5). The product of the single substitution frequencies in the controls is nonetheless strongly correlated with the double frequency (Pearson correlation coefficients r=0.93 for syn33 and r=0.76 for syn31). A significant correlation was also observed between the single and double frequencies in all four classes of double substitutions (Fig. S6). We further verified that, for the NS cases, the observed frequencies of double substitutions were significantly higher than expected from the frequencies of single substitutions, with the addition of simultaneous double mutations rates estimated from the controls (paired t-test p-val=6.9×10^-04^ and signed rank test p-val=0.0011). This result presents further evidence that, although simultaneous double mutations contribute to the observed double substitution frequency and to the DF, they cannot account for the elevated values in NS. Thus, the increase in DF in these cases can only be attributed to positive selection.

## Discussion

The central goal of this work was to comprehensively characterize the selective landscape of codon double substitutions by accurately taking account the mutational biases in the inference of selection. The control for mutation biases was achieved by comparing the DF for codon double substitutions to those of double synonymous substitutions. Previously analyzed codon double substitutions in serine codons^17^ and in stop codons^16^ suggested that these changes are under positive selection due to elevated double substitution frequencies compare to the expectation from single substitutions. Our focus here was to infer the type of selection by using more adequate controls, namely equivalent synonymous double substitutions, in order to address the possibility that apparent selection affecting codon double substitutions was due to mutational biases as previously suggested^9,14^. Indeed, we observed that adjacent double synonymous substitutions (syn_31) had a higher DF compared to the corresponding non-adjacent substitutions (syn_33), although this difference was not statistically significant (Fig. 1E).

Partitioning of codon double substitutions into 4 classes based on the (non)synonymy of the intermediate and final codon to the ancestral codon (SS, SN, NS and NN) predicts the type of selection affecting the second step of the respective double substitutions (Fig. 1). In fact, this classification is a simple derivative of the classification of single substitutions in protein coding genes into synonymous substitutions that are generally assumed to evolve neutrally, and non-synonymous substitutions most of which are subject to purifying selection^20^ (Fig. S1). The classes of double substitutions are the four possible combinations of synonymous and non-synonymous substitutions at each step. Because the state resulting from the second step is the one that is fixed during evolution, the nature of this step largely defines the selective regime of the double substitution perceived as one evolutionary event (Fig. 2A). Thus, SS doubles are effectively neutral. The SN doubles that drive an amino acid site away from the original state are generally subject to purifying selection, the strength of which depends on the similarity between the new amino acid introduced by the second substitution and the original amino acid. The few SN cases that appear to be driven by positive selection all involve conservative amino acid replacements and might reflect a hitherto unrecognized process of adaptive fine-tuning of protein structures. Alternatively, this apparent positive selection could be an artifact caused context-specific mutational biases. The NS doubles that return the site to the ancestral state are positively selected because, by definition, in all these cases, the similarity of the final (same as ancestral) amino acid to the ancestral one is always greater compared to the intermediate. The NN doubles are heterogeneous, evolving either under purifying selection or under positive selection depending on which amino acid, intermediate or final, is more similar to the ancestral one. Notably, the DAS values are not always positive for the NN cases under positive selection, as generally expected. This is most likely due to the fact that each amino acid substitution matrix accurately reflects similarity in certain properties but not others, and thus, does not equally well apply to all amino acid replacements. No single matrix is expected to be fully compatible with selection regimes on codon substitutions because they represent a mixture of numerous proteins from many environments that are subject to different sets of functional constraints.

Overall, the results of the present, comprehensive analysis of the evolutionary regimes of double substitutions reaffirm the predominantly conservative character of protein evolution^5,40^. In bulk, all classes of double substitutions can be viewed as evolving under purifying selection if the double is taken as one evolutionary event. The positive selection detected for the second steps of the NS and many NN doubles is a consequence of the deleterious effect of the first substitution. The conclusion on the overall dominance of purifying selection is further supported by the comparison of double substitutions in fast vs. slow evolving genes. In accord with the identified purifying selection on SN cases, these have significantly greater DF in fast evolving genes, similar to the higher rate of single non-synonymous changes in fast evolving genes compared to slow evolving ones. Conversely, those NS and NN substitutions, for which the second step was found to be driven by positive selection, showed a higher DF in slow evolving genes.

Compensation for the effects of deleterious mutations through subsequent positive selection has been previously hypothesized and demonstrated in other evolutionary contexts^41-43^. A major implication of the present results is that fitness valleys are commonly crossed in codon evolution as a result of positive selection that follows a deleterious non-synonymous mutation and that this route of evolution is, in large part, determined by the organization of the genetic code itself.

## Materials and methods

### Datasets

Genomic data for bacteria and archaea were obtained from an updated version of the ATGC (Alignable Tight Genome Clusters) database^27^. To reconstruct the history of nucleotide substitutions in protein-coding DNA under the parsimony principle, we used triplets of closely related species as previously described^16,17,44^. Alignments of all sequences in each ATGC COG (Cluster of Orthologous Genes) were constructed using the MAFFT software with the -linsi parameter^45^. The genes were divided into slow and fast evolving ones by comparing the dN/dS value of each gene to the median dN/dS among all genome triplets in the given ATGC.

### Analysis of codon double substitutions

For each codon change, the frequency of change to any other codon was calculated as the number of such changes divided by the number of ancestral reconstructions of the given codon. For each double substitution, the double fraction (DF) was calculated as the observed double substitution frequency divided by the cumulative single substitution frequency plus the double frequency. For example, for the change AAA→GGA, the DF was the observed frequency of AAA→GGA divided by the cumulative counts of AAA→GAA and AAA→AGA and AAA→GGA (under the assumption that the double substitution occurred as a result of two consecutive single substitutions). Thus, for each double substitution, the following values were collected and estimated:

1. The double substitution count.
2. The cumulative single substitution count (which is the sum of the two single counts and the double count).
3. The ancestral state count – count of all cases where the originating codon is inferred as ancestral under the parsimony principal.
4. double substitution frequency – ‘double substitution count’ / ‘ancestral state count’.
5. single cumulative frequency – ‘single substitution count’ / ‘ancestral state count’.
6. double fraction (DF) – double substitution frequency divided by cumulative single frequency – equivalent to the double substitution count divided by the cumulative single substitution count.

### Assignment of codon double substitution types

For each codon double substitution, there are two distinct paths from the ancestral codon state to the final (derived) codon state, where each step in the path is a single substitution to or from an intermediate codon state. Each step can be either synonymous or non-synonymous, and the ancestral vs. final codon also can be either synonymous or non-synonymous. Some codon substitutions include a stop codon as the intermediate in one of the paths; these cases were disregarded in the current analysis. Each codon double substitution was assigned one of the 4 combination types based on the (non)synonymy of the ancestral to the intermediate codons, and the (non)synonymy of the ancestral vs. the final codon state. The 4 classes are as follows: -

1. SS, codon double substitutions in which both intermediates and the final codon are all synonymous
2. SN, codon double substitutions in which at least one intermediate is synonymous whereas the final codon is non-synonymous to the ancestral codon
3. NS, codon double substitutions in which one of the intermediates in non-synonymous whereas the final codon is synonymous to the ancestral codon
4. NN, codon double substitutions in which both intermediates are non-synonymous, and the final codon is also non-synonymous to the ancestral one.

### Analysis of double synonymous substitutions in adjacent codons: the null models

For double synonymous substitutions in adjacent codons, we collected the same data as for the codon double substitutions, in codon-like 3-base sequences with 3 configurations:

A. A constant 2^nd^ codon positions followed by a 4-fold degenerate site in the 3^rd^ codon positions which is followed by a 2-fold degenerate site in the 1^st^ codon position of the next codon (Fig. 1A).
B. A 4-fold degenerate site in the 3^rd^ codon positions which is followed by a 2-fold degenerate site in the 1^st^ codon position of the next codon, which is followed by a constant base in the 2^nd^ codon position of the second codon (Fig. 1B).
C. A 4-fold degenerate site in the 3^rd^ codon positions which is followed by a constant 1^st^ codon position in the second codon of which the 2^nd^ position is disregarded and followed by a 4-fold degenerate site in the 3^rd^ codon position (Fig. 1C).

The first codon in configurations A and B can be any of the 4-fold degenerate codons, i.e, codons for L, V, S, P Y, A, R and G, and the second codon in these configurations can be either a codon for either R or L which are the only two amino acids that have a degenerate 1^st^ codon position. An additional restriction for configurations A and B is that the ancestral state of the 3^rd^ codon position of the 2^nd^ codon is a purine (A/G) because only then can the 1^st^ codon substitution be synonymous. The 1^st^ and 2^nd^ codons of configuration C can be any of the 4-fold degenerate codons.

### Analysis of double substitutions in non-coding intergenic regions

Codon double substitutions were also compared to double substitutions in non-coding intergenic regions. The same analysis was performed on all possible frames of the aligned non-coding sequences as for the coding genes, treating base triplets of bases as codons.

### Estimation of simultaneous double mutation frequency

In the null models syn31 and syn33, the expected frequency of double substitutions, in the absence of simultaneous double mutations, can be estimated by the product of the frequencies of single synonymous substitutions. Because both single substitutions are synonymous, the effect of their order is assumed to be minimal. Thus, the estimated contribution of simultaneous double mutations in these cases is the difference between the observed double substitution frequency and the product of the corresponding single substitutions. In 12 of the 634 cases, the observed double frequency was smaller than the product of single substitution frequencies; these cases were ignored. We further estimated the expected rate of NS substitutions, under the assumption of neutrality at the second step, as the weighted mean of the products of each of the corresponding single substitution frequencies multiplied by the equivalent null model’s synonymous frequency which is expected if the second step is neutral. To each expected NS value, the estimated simultaneous double mutational rate was added. If the observed frequency of NS double substitutions can be explained by simultaneous double mutations, then the expected rate plus the double mutational rate should be equal to the observed NS frequency. Thus, to assess the contribution of selection, the expected frequency (after adding the double mutation frequency) was subtracted from the observed double substitution frequency.

### Statistical tests

Two samples t-test and the non-parametric Wilcoxon Ranksum test were used to compare the DF values between each of the codon double substitution types (SS, SN, NS, NN) and each of the null models (syn-31, syn-33) and between each of the codon double substitution types and adjacent and non-adjacent double substitutions in non-coding intergenic regions. Alpha level for significance was 0.01.

Paired t-test and signed rank test were used to compare between the DF of different codon double substitution types in fast vs. slow evolving genes. Alpha level for significance was 0.01. Fisher’s exact test was used to compare the number of double codon substitutions to single cumulative substitutions, to test for significant differences in the DF between a specific codon double substitution and the comparable null model. For example, the codon double substitution GCC→GTA, which changes the encoded amino acid from A to V, is compared to the null model of two adjacent synonymous substitutions with configuration A (Fig. 1A) where the 1^st^ base is G in the 2^nd^ codon position, followed by a synonymous C→T change in a 4-fold degenerate 3^rd^ codon position and by a synonymous C→A change in the 1^st^ codon position of the next codon (coding for R). An example of the comparison for the non-adjacent codon double substitution CTT→TTA is detailed in Fig. 1D. The Benjamini–Hochberg procedure was used to correct for multiple testing, with alpha of 0.05.

