## Supplementary information for "Crossing fitness valleys via double substitutions within codons"

### Evidence for positive selection acting on codon double substitutions in prokaryotes

Table S1 – Individual comparison of NN codon double substitution to a single null model with the same mutational changes, and adjacency (syn_31 for adjacent substitutions and syn_33 for non-adjacent substitutions) using Fisher's exact test

| codon change | | Null model - double syn | | | Codon change | | | Fisher double vs. single | Amino acid change | | | | mode of selection after BH |
|---|---|---|---|---|---|---|---|---|---|---|---|---|---|
| ancestral | derived | double | single | ancestral | double | single | ancestral | | ancestral | derived | inter-1 | inter-2 | |
| CAG | AAC | 46 | 5089 | 72658 | 226 | 4185 | 630066 | 3.99E-37 | Q | N | H | K | >> |
| TTC | CTG | 89 | 1853 | 16845 | 528 | 3052 | 590362 | 5.02E-34 | F | L | L | L | >> |
| TTT | CTG | 44 | 1759 | 25416 | 257 | 2983 | 386236 | 4.7E-17 | F | L | L | L | >> |
| TTA | ATT | 58 | 1227 | 22010 | 330 | 2355 | 269701 | 3.35E-16 | L | I | F | I | >> |
| CAG | GAT | 38 | 2868 | 72658 | 208 | 4460 | 630066 | 1.72E-15 | Q | D | H | E | >> |
| CAC | GAA | 74 | 4556 | 119063 | 62 | 1003 | 267623 | 3.47E-13 | H | E | Q | D | >> |
| TTC | ATG | 22 | 637 | 16845 | 192 | 1325 | 590362 | 3.8E-13 | F | M | L | I | >> |
| CAC | AAG | 78 | 5948 | 119063 | 82 | 1940 | 267623 | 8.99E-13 | H | K | Q | N | >> |
| TTG | ATT | 14 | 621 | 16226 | 196 | 1705 | 204259 | 9.38E-13 | L | I | F | M | >> |
| AAG | GCG | 42 | 1149 | 20988 | 296 | 2788 | 534568 | 1.94E-12 | K | A | T | E | >> |
| ATA | CTG | 14 | 892 | 10829 | 169 | 2425 | 146332 | 1.86E-10 | I | L | M | L | >> |
| TGG | GGC | 115 | 3048 | 51165 | 46 | 358 | 350601 | 6.71E-10 | W | G | C | G | >> |
| TTT | ATG | 20 | 951 | 25416 | 146 | 1894 | 386236 | 1.09E-09 | F | M | L | I | >> |
| ATT | CTG | 5 | 463 | 11031 | 338 | 4306 | 474289 | 1.53E-09 | I | L | M | L | >> |
| GAG | AAT | 35 | 2623 | 70692 | 227 | 6505 | 668983 | 7.95E-09 | E | N | D | K | >> |
| CAG | AAT | 12 | 1059 | 72658 | 178 | 3801 | 630066 | 1.22E-08 | Q | N | H | K | >> |
| ATA | GTG | 45 | 1176 | 10829 | 506 | 5959 | 146332 | 4.23E-08 | I | V | M | V | >> |
| GAC | CAA | 49 | 4127 | 80232 | 130 | 4726 | 700855 | 2.01E-07 | D | Q | E | H | >> |
| ATC | CTG | 37 | 530 | 7969 | 905 | 5754 | 752044 | 3.30E-07 | I | L | M | L | >> |
| CCG | GAG | 32 | 1876 | 45403 | 165 | 3851 | 580630 | 3.50E-07 | P | E | Q | A | >> |
| ACG | CAG | 16 | 378 | 9609 | 166 | 1193 | 340066 | 4.98E-07 | T | Q | K | P | >> |
| AAC | CAG | 31 | 771 | 11330 | 201 | 2010 | 436160 | 5.95E-07 | N | Q | K | H | >> |
| GCG | CAG | 125 | 2103 | 79948 | 443 | 4544 | 907397 | 0.00000103 | A | Q | E | P | >> |
| TTC | GTG | 35 | 839 | 16845 | 141 | 1386 | 590362 | 0.00000124 | F | V | L | V | >> |
| ATT | TTG | 17 | 592 | 11031 | 236 | 2748 | 474289 | 0.00000131 | I | L | M | F | >> |
| ATC | GTG | 38 | 777 | 7969 | 1549 | 15330 | 752044 | 0.00000226 | I | V | M | V | >> |
| TTG | ATC | 19 | 560 | 16226 | 136 | 1366 | 204259 | 0.00000239 | L | I | F | M | >> |
| TTA | ATC | 29 | 907 | 22010 | 149 | 1889 | 269701 | 0.00000262 | L | I | F | I | >> |
| TTA | GTT | 24 | 1085 | 22010 | 113 | 1919 | 269701 | 0.0000038 | L | V | F | V | >> |
| AAG | GAT | 31 | 1195 | 12030 | 154 | 2512 | 534568 | 0.00000458 | K | D | N | E | >> |
| CAG | ACG | 20 | 740 | 14713 | 252 | 3516 | 630066 | 0.00000498 | Q | T | P | K | >> |
| TGG | CGC | 346 | 7482 | 51165 | 51 | 521 | 350601 | 0.00000852 | W | R | C | R | >> |
| ATC | TTG | 9 | 349 | 7969 | 204 | 2178 | 752044 | 0.0000113 | I | L | M | F | >> |
| TTT | ATA | 48 | 1277 | 25416 | 138 | 1788 | 386236 | 0.0000148 | F | I | L | I | >> |
| CAC | AAA | 25 | 1717 | 119063 | 58 | 1487 | 267623 | 0.0000283 | H | K | Q | N | >> |
| ATG | CTT | 11 | 568 | 9430 | 290 | 4864 | 642679 | 0.0000305 | M | L | I | L | >> |
| ATT | GTG | 22 | 795 | 11031 | 744 | 11951 | 474289 | 0.0000446 | I | V | M | V | >> |
| ATA | TTG | 30 | 949 | 10829 | 179 | 2793 | 146332 | 0.000198 | I | L | M | L | >> |
| TTT | GTA | 30 | 1196 | 25416 | 92 | 1724 | 386236 | 0.000308 | F | V | L | V | >> |
| TTG | GTT | 21 | 995 | 16226 | 70 | 1393 | 204259 | 0.000309 | L | V | F | V | >> |
| CAA | CGC | 24 | 672 | 12911 | 184 | 2490 | 270512 | 0.000525 | Q | R | H | R | >> |
| ATG | GTC | 46 | 1463 | 9430 | 229 | 4236 | 642679 | 0.000627 | M | V | I | V | >> |
| ACT | GTT | 353 | 5747 | 27743 | 215 | 2569 | 117239 | 0.000638 | T | V | I | A | >> |
| CAG | GCG | 45 | 700 | 14713 | 469 | 4362 | 630066 | 0.00105 | Q | A | P | E | >> |
| ATG | GTT | 36 | 1414 | 9430 | 176 | 3885 | 642679 | 0.0014 | M | V | I | V | >> |
| AAC | CAA | 17 | 764 | 11330 | 109 | 2228 | 436160 | 0.00156 | N | Q | K | H | >> |
| TGG | GGT | 98 | 3191 | 51165 | 23 | 338 | 350601 | 0.00168 | W | G | C | G | >> |
| AAT | CAG | 14 | 676 | 14264 | 150 | 3176 | 333464 | 0.00201 | N | Q | K | H | >> |
| AGT | GGG | 154 | 5198 | 56365 | 61 | 1254 | 114762 | 0.00217 | S | G | R | G | >> |
| ATG | TTA | 35 | 819 | 9430 | 409 | 5609 | 642679 | 0.00221 | M | L | I | L | >> |
| TTT | CTA | 51 | 2064 | 25416 | 114 | 2742 | 386236 | 0.00226 | F | L | L | L | >> |
| CAC | CGA | 19 | 2948 | 20901 | 41 | 2743 | 267623 | 0.00253 | H | R | Q | R | >> |
| ATG | CTC | 24 | 620 | 9430 | 377 | 5249 | 642679 | 0.00279 | M | L | I | L | >> |
| AAC | GAA | 37 | 1177 | 11330 | 199 | 3737 | 436160 | 0.0028 | N | E | K | D | >> |
| CAG | GAC | 281 | 7107 | 72658 | 247 | 4835 | 630066 | 0.00433 | Q | D | H | E | >> |

| | | | | | | | | | | | | | |
|---|---|---|---|---|---|---|---|---|---|---|---|---|---|
| TTA | GTC | 11 | 781 | 22010 | 54 | 1575 | 269701 | 0.00478 | L | V | F | V | >> |
| AAG | GAC | 49 | 1309 | 12030 | 134 | 2231 | 534568 | 0.0057 | K | D | N | E | >> |
| AGT | GGA | 200 | 5605 | 56365 | 70 | 1313 | 114762 | 0.0058 | S | G | R | G | >> |
| GAC | GGA | 4 | 953 | 8102 | 116 | 7884 | 700855 | 0.00658 | D | G | E | G | >> |
| TGG | AGT | 109 | 2410 | 51165 | 19 | 203 | 350601 | 0.00743 | W | S | C | R | >> |
| AAT | GAA | 55 | 1413 | 14264 | 367 | 6453 | 333464 | 0.00878 | N | E | K | D | >> |
| TGG | AGC | 75 | 2216 | 51165 | 15 | 196 | 350601 | 0.0101 | W | S | C | R | >> |
| TGG | CGT | 215 | 7511 | 51165 | 25 | 498 | 350601 | 0.0145 | W | R | C | R | >> |
| TTC | ATA | 14 | 604 | 16845 | 46 | 954 | 590362 | 0.0149 | F | I | L | I | >> |
| AGA | GGC | 106 | 4750 | 54066 | 15 | 328 | 94676 | 0.015 | R | G | S | G | >> |
| GAG | ACG | 21 | 827 | 8129 | 243 | 5614 | 668983 | 0.0179 | E | T | A | K | >> |
| TTG | GTC | 25 | 933 | 16226 | 53 | 1097 | 204259 | 0.0199 | L | V | F | V | >> |
| CAC | GAG | 389 | 9049 | 119063 | 84 | 1454 | 267623 | 0.0211 | H | E | Q | D | >> |
| AAT | CAA | 32 | 912 | 14264 | 211 | 3878 | 333464 | 0.0227 | N | Q | K | H | >> |
| TTT | GTG | 25 | 893 | 25416 | 84 | 1814 | 386236 | 0.0286 | F | V | L | V | >> |
| AGG | GGC | 128 | 3383 | 29349 | 18 | 265 | 36831 | 0.0345 | R | G | S | G | >> |
| TTC | TCA | 8 | 469 | 10933 | 30 | 756 | 590362 | 0.0401 | F | S | L | S | >> |
| GTT | TCT | 94 | 5428 | 38436 | 48 | 1915 | 250979 | 0.0429 | V | S | A | F | = |
| AGC | CGA | 65 | 2201 | 34345 | 25 | 1379 | 344526 | 0.0475 | S | R | R | R | = |
| AGA | GGT | 170 | 5169 | 54066 | 18 | 327 | 94676 | 0.0596 | R | G | S | G | = |
| CCT | ATT | 150 | 5975 | 136206 | 21 | 528 | 122662 | 0.0648 | P | I | L | T | = |
| GAG | CCG | 13 | 359 | 8129 | 134 | 6398 | 668983 | 0.0651 | E | P | A | Q | = |
| ATG | CTA | 44 | 1043 | 9430 | 218 | 7065 | 642679 | 0.0752 | M | L | I | L | = |
| AGC | TGG | 65 | 1799 | 34345 | 15 | 697 | 344526 | 0.0757 | S | W | R | C | = |
| CAA | AAC | 23 | 487 | 7534 | 121 | 3915 | 270512 | 0.0795 | Q | N | H | K | = |
| AGC | GGA | 122 | 3719 | 34345 | 128 | 4881 | 344526 | 0.0922 | S | G | R | G | = |
| GAC | CAG | 301 | 6756 | 80232 | 274 | 7085 | 700855 | 0.0971 | D | Q | E | H | = |
| GAC | GTA | 2 | 179 | 8102 | 12 | 4280 | 700855 | 0.108 | D | V | E | V | = |
| AAC | GAG | 64 | 1197 | 11330 | 141 | 3369 | 436160 | 0.124 | N | E | K | D | = |
| GAG | CAT | 40 | 3090 | 70692 | 69 | 7240 | 668983 | 0.141 | E | H | D | Q | = |
| AGC | CGG | 70 | 1960 | 34345 | 62 | 1332 | 344526 | 0.148 | S | R | R | R | = |
| ATG | TTC | 14 | 395 | 9430 | 187 | 3412 | 642679 | 0.15 | M | F | I | L | = |
| AGT | CGG | 41 | 2503 | 56365 | 11 | 396 | 114762 | 0.151 | S | R | R | R | = |
| CAA | GAT | 14 | 373 | 7534 | 162 | 2863 | 270512 | 0.178 | Q | D | H | E | = |
| CAA | AAT | 21 | 463 | 7534 | 236 | 3728 | 270512 | 0.179 | Q | N | H | K | = |
| AAA | GAT | 74 | 1884 | 20660 | 273 | 5775 | 574412 | 0.18 | K | D | N | E | = |
| CCG | AAG | 36 | 987 | 45403 | 90 | 1874 | 580630 | 0.18 | P | K | Q | T | = |
| GAC | GCA | 9 | 276 | 8102 | 114 | 5656 | 700855 | 0.19 | D | A | E | A | = |
| CTT | GCT | 305 | 2963 | 30749 | 67 | 786 | 228202 | 0.202 | L | A | P | V | = |
| TTT | ACT | 166 | 3584 | 78483 | 41 | 1113 | 386236 | 0.21 | F | T | S | I | = |
| AAT | GAG | 36 | 1176 | 14264 | 219 | 5664 | 333464 | 0.235 | N | E | K | D | = |
| TGT | GGG | 174 | 4761 | 100899 | 8 | 152 | 60699 | 0.281 | C | G | W | G | = |
| TGC | CGG | 463 | 13010 | 84580 | 16 | 343 | 154212 | 0.303 | C | R | W | R | = |
| TCT | ATT | 90 | 2307 | 31431 | 44 | 1365 | 134939 | 0.318 | S | I | F | T | = |
| TTT | GCT | 233 | 3461 | 78483 | 66 | 1120 | 386236 | 0.367 | F | A | S | V | = |
| TGT | AGG | 164 | 5460 | 100899 | 3 | 187 | 60699 | 0.377 | C | R | W | S | = |
| CAA | CCC | 9 | 317 | 12911 | 32 | 1628 | 270512 | 0.391 | Q | P | H | P | = |
| TGT | CGG | 228 | 8621 | 100899 | 5 | 129 | 60699 | 0.401 | C | R | W | R | = |
| ATG | GTA | 78 | 1898 | 9430 | 290 | 6272 | 642679 | 0.411 | M | V | I | V | = |
| GAA | GGC | 9 | 254 | 4630 | 217 | 7695 | 747383 | 0.445 | E | G | D | G | = |
| TTC | GTA | 17 | 796 | 16845 | 29 | 1049 | 590362 | 0.453 | F | V | L | V | = |
| CAC | CCA | 5 | 616 | 20901 | 11 | 863 | 267623 | 0.455 | H | P | Q | P | = |
| AGG | TGC | 58 | 1390 | 29349 | 3 | 126 | 36831 | 0.476 | R | C | S | W | = |
| ACT | CTT | 109 | 2255 | 27743 | 44 | 1050 | 117239 | 0.477 | T | L | I | P | = |
| AAG | CCG | 18 | 590 | 20988 | 92 | 3640 | 534568 | 0.486 | K | P | T | Q | = |
| AAA | CAT | 29 | 1109 | 20660 | 139 | 6043 | 574412 | 0.519 | K | H | N | Q | = |
| ATG | TTT | 18 | 360 | 9430 | 185 | 3112 | 642679 | 0.554 | M | F | I | L | = |
| CAT | GAG | 22 | 466 | 8000 | 59 | 1447 | 215209 | 0.598 | H | E | Q | D | = |
| GAT | CAG | 14 | 461 | 8454 | 195 | 5310 | 558773 | 0.602 | D | Q | E | H | = |
| GAA | AAT | 66 | 1321 | 11884 | 503 | 10709 | 747383 | 0.631 | E | N | D | K | = |
| CAA | GAC | 13 | 394 | 7534 | 91 | 3094 | 270512 | 0.64 | Q | D | H | E | = |
| CAT | AAG | 13 | 448 | 8000 | 76 | 2192 | 215209 | 0.666 | H | K | Q | N | = |
| CCT | GTT | 397 | 10755 | 136206 | 27 | 679 | 122662 | 0.676 | P | V | L | A | = |

| | | | | | | | | | | | | | |
|---|---|---|---|---|---|---|---|---|---|---|---|---|---|
| CAT | AAA | 15 | 339 | 8000 | 103 | 1993 | 215209 | 0.687 | H | K | Q | N | = |
| AGG | GGT | 140 | 3520 | 29349 | 11 | 252 | 36831 | 0.739 | R | G | S | G | = |
| GAC | AAA | 36 | 2751 | 80232 | 112 | 7939 | 700855 | 0.776 | D | K | E | N | = |
| AAG | CAT | 16 | 580 | 12030 | 90 | 3504 | 534568 | 0.778 | K | H | N | Q | = |
| ACG | GAG | 27 | 1076 | 9609 | 171 | 6313 | 340066 | 0.838 | T | E | K | A | = |
| CAA | CTC | 8 | 404 | 12911 | 33 | 1414 | 270512 | 0.849 | Q | L | H | L | = |
| AAA | GAC | 42 | 1572 | 20660 | 134 | 5205 | 574412 | 0.856 | K | D | N | E | = |
| AGT | TGG | 132 | 3959 | 56365 | 9 | 307 | 114762 | 0.868 | S | W | R | C | = |
| CAT | GAA | 20 | 353 | 8000 | 74 | 1236 | 215209 | 0.899 | H | E | Q | D | = |
| TTC | CTA | 42 | 1781 | 16845 | 58 | 2357 | 590362 | 0.919 | F | L | L | L | = |
| GAT | CAA | 10 | 343 | 8454 | 157 | 4950 | 558773 | 1 | D | Q | E | H | = |
| CAC | CTA | 3 | 428 | 20901 | 6 | 844 | 267623 | 1 | H | L | Q | L | = |
| AGT | CGA | 67 | 2890 | 56365 | 10 | 445 | 114762 | 1 | S | R | R | R | = |
| TCT | GTT | 125 | 2874 | 31431 | 45 | 2031 | 134939 | 0.0000908 | S | V | F | A | << |
| GCT | ATT | 1833 | 32582 | 417377 | 186 | 4440 | 232600 | 0.000115 | A | I | V | T | << |
| ACT | TTT | 102 | 1989 | 27743 | 37 | 1453 | 117239 | 0.000212 | T | F | I | S | << |
| GAA | CAT | 19 | 522 | 11884 | 107 | 7891 | 747383 | 0.00034 | E | H | D | Q | << |
| CTT | ACT | 96 | 2792 | 30749 | 42 | 2305 | 228202 | 0.000502 | L | T | P | I | << |
| GAC | AAG | 114 | 5206 | 80232 | 142 | 10184 | 700855 | 0.000516 | D | K | E | N | << |
| TCT | CTT | 273 | 4820 | 31431 | 41 | 1274 | 134939 | 0.000557 | S | L | F | P | << |
| GCT | CTT | 952 | 23864 | 417377 | 55 | 2171 | 232600 | 0.000837 | A | L | V | P | << |
| AAG | CAC | 41 | 701 | 12030 | 101 | 3254 | 534568 | 0.00124 | K | H | N | Q | << |
| ATT | TCT | 90 | 2785 | 44754 | 30 | 1691 | 474289 | 0.00403 | I | S | T | F | << |
| GAG | AAC | 162 | 5662 | 70692 | 162 | 7829 | 668983 | 0.00437 | E | N | D | K | << |
| GAA | GTC | 6 | 167 | 4630 | 55 | 6177 | 747383 | 0.00578 | E | V | D | V | << |
| AGC | GGG | 135 | 3486 | 34345 | 133 | 4802 | 344526 | 0.00822 | S | G | R | G | << |
| GCT | TTT | 477 | 18909 | 417377 | 63 | 3540 | 232600 | 0.00836 | A | F | V | S | << |
| GAA | AAC | 51 | 1314 | 11884 | 297 | 11359 | 747383 | 0.0128 | E | N | D | K | << |
| TGC | AGG | 94 | 2993 | 84580 | 4 | 398 | 154212 | 0.0153 | C | R | W | S | << |
| GCG | AAG | 99 | 2973 | 79948 | 278 | 11089 | 907397 | 0.0183 | A | K | E | T | << |
| TGC | GGG | 255 | 4347 | 84580 | 8 | 307 | 154212 | 0.0192 | C | G | W | G | << |
| AAA | CAC | 22 | 822 | 20660 | 88 | 5561 | 574412 | 0.0321 | K | H | N | Q | << |
| ATT | GCT | 524 | 4640 | 44754 | 182 | 10538 | 474289 | 2.06E-119 | I | A | T | V | << |
| GAG | CAC | 342 | 6304 | 70692 | 102 | 8662 | 668983 | 3.23E-49 | E | H | D | Q | << |
| TTT | CCT | 336 | 5448 | 78483 | 33 | 2083 | 386236 | 2.92E-18 | F | P | S | L | << |
| ATT | CCT | 99 | 2502 | 44754 | 27 | 3144 | 474289 | 1.14E-14 | I | P | T | L | << |
| GAA | CAC | 36 | 547 | 11884 | 98 | 8738 | 747383 | 2.05E-14 | E | H | D | Q | << |
| GTT | ACT | 348 | 7731 | 38436 | 239 | 9770 | 250979 | 6.9E-13 | V | T | A | I | << |
| GTT | CCT | 199 | 5543 | 38436 | 18 | 2026 | 250979 | 2E-11 | V | P | A | L | << |
| CTT | TCT | 255 | 5036 | 30749 | 39 | 2045 | 228202 | 7.51E-10 | L | S | P | F | << |
| GAT | AAG | 46 | 982 | 8454 | 172 | 11659 | 558773 | 7.67E-10 | D | K | E | N | << |
| CCT | TTT | 656 | 10829 | 136206 | 38 | 1458 | 122662 | 3.03E-08 | P | F | L | S | << |
| GAT | AAA | 46 | 868 | 8454 | 277 | 11442 | 558773 | 0.00000943 | D | K | E | N | << |

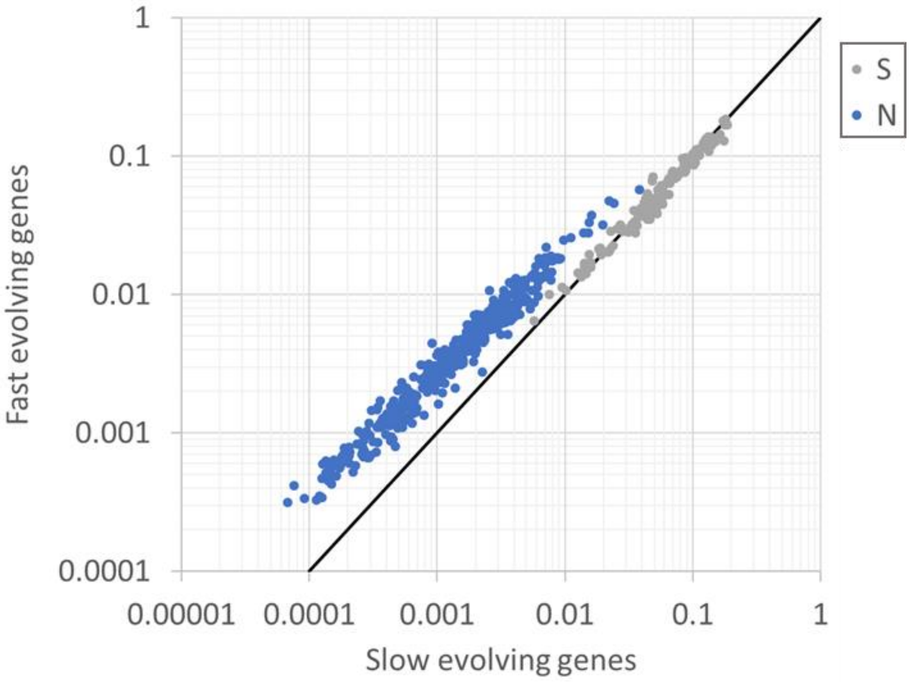

**Figure S1.** Comparison of single synonymous (S) and non-synonymous (N) substitution frequencies in fast evolving genes (with dN/dS>median) vs. frequencies in slow evolving genes (dN/dS<media).

A

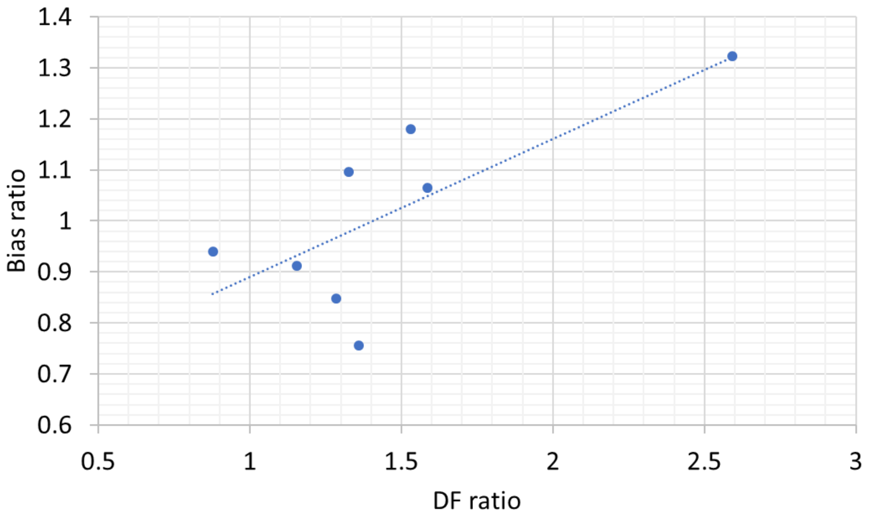

B

| Codon | Amino acid | Relative frequency in fast genes | Relative frequency in slow genes | Relative bias | SS case | Bias ratio |
|---|---|---|---|---|---|---|
| AGA | R | 0.218748 | 0.272035 | 1.243602 | CGG->AGA | 1.3220813 |
| AGG | R | 0.100178 | 0.093699 | 0.93533 | CGA->AGG | 1.0641132 |
| CGA | R | 0.103466 | 0.090944 | 0.878976 | AGG->CGA | 0.9397496 |
| CGG | R | 0.577609 | 0.543322 | 0.94064 | AGA->CGG | 0.7563831 |
| CTA | L | 0.034539 | 0.027896 | 0.807679 | TTG->CTA | 0.9119221 |
| CTG | L | 0.664403 | 0.70345 | 1.058769 | TTA->CTG | 1.1795942 |
| TTA | L | 0.169185 | 0.151856 | 0.897571 | CTG->TTA | 0.8477492 |
| TTG | L | 0.131872 | 0.116798 | 0.885688 | CTA->TTG | 1.0965849 |

**Figure S2.** (A) Pearson correlation (R=0.74, p-val=0.037) between selection strength in the SS class, estimated by the DF ratio (ratio between the individual codon DFs and the control DF-individual cases of syn 33 as detailed in Fig. 1D), and the codon bias ratio. (B) The bias ratio is calculated as the ratio between the biases of the final codon compared to the original one, based on relative biases of these codon calculated as the ratio of their relative frequency in slow vs. fast evolving genes (see methods for detailed explanation on fast and slow genes).

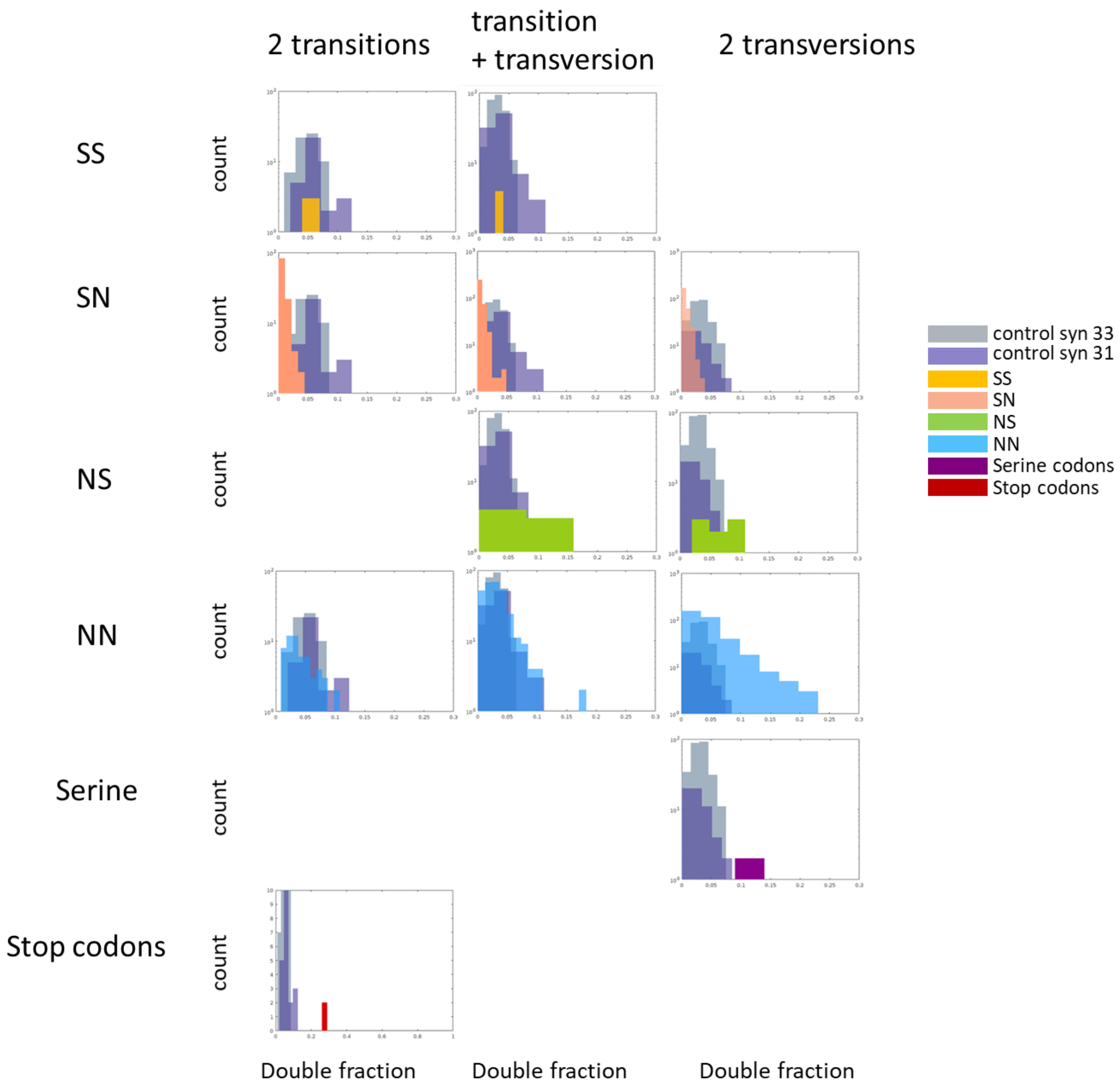

**Figure S3.** Comparison of each of the codon double substitution classes to the double synonymous null models, separately by transitions and transversions.

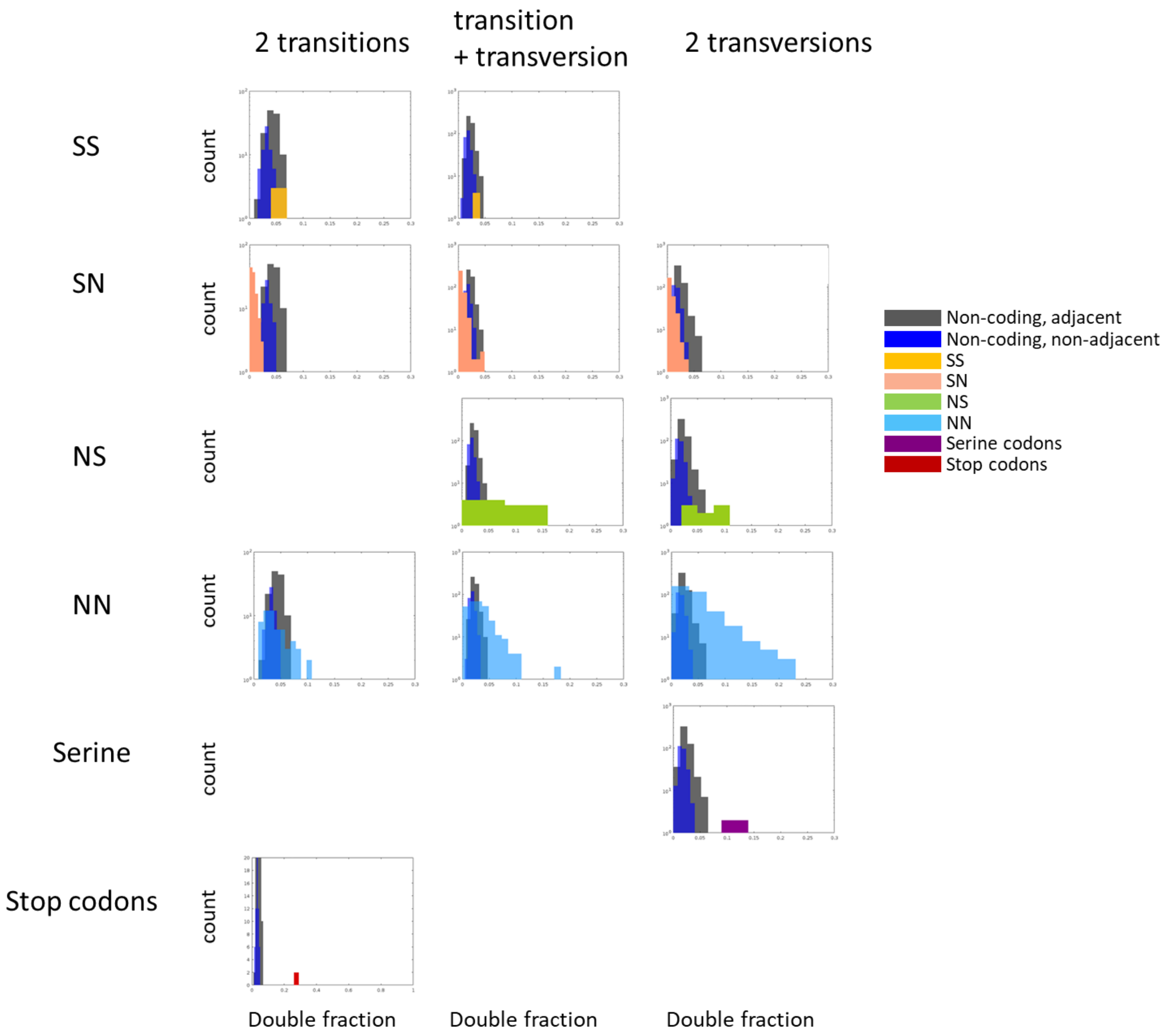

**Figure S4.** Comparison of each of the codon double substitution classes to non-coding codon-like base triplets, separately by transitions and transversions.

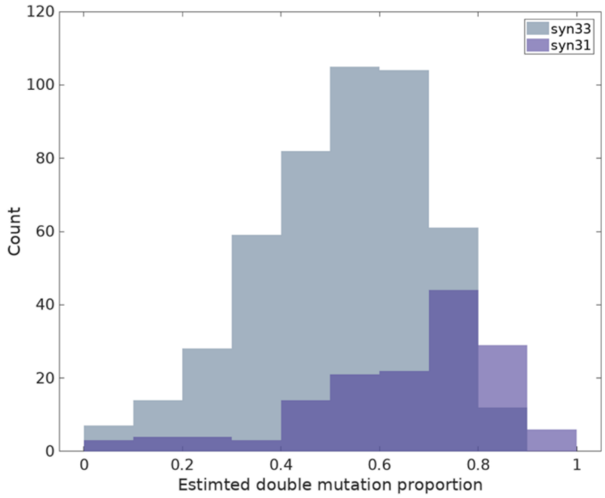

**Figure S5.** Proportion of the estimated double mutation frequency out of the observed double substitution frequency in the null models syn31 (adjacent synonymous substitutions) and syn33 (non-adjacent synonymous substitutions). The difference between the syn31 and syn33 distributions is significant (p-val= $9.98 \times 10^{-15}$ for t-test and p-val= $2.02 \times 10^{-16}$ for rank sum test).

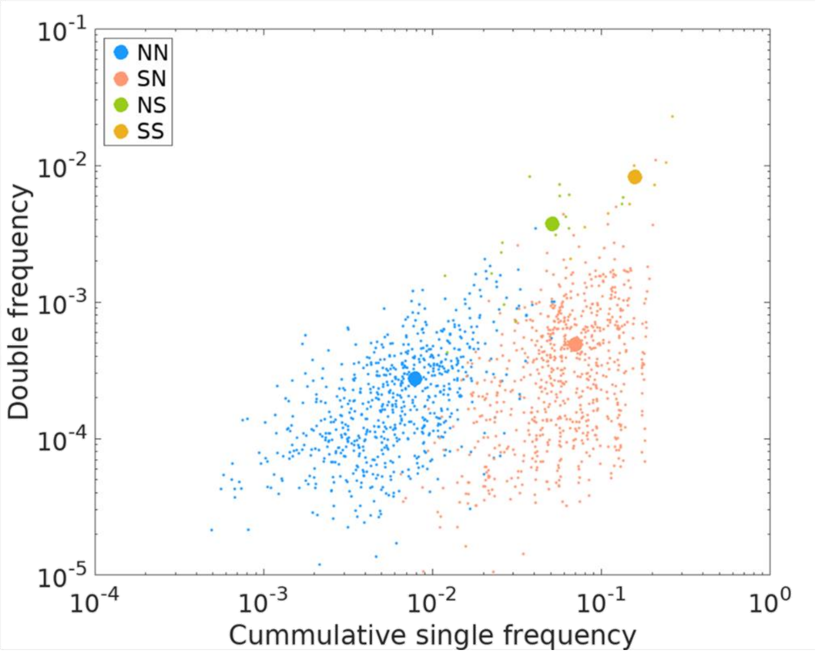

**Figure S6.** Dependency of double frequency on the cumulative single frequency (Spearman correlation of: 0.56, 0.4, 0.97 and 0.7 for NN, SN, SS and NS respectively, with respective p-values: $6.018 \times 10^{-58}$, $3.15 \times 10^{-29}$, $3.96 \times 10^{-04}$ and $3.23 \times 10^{-03}$), and the separation of the four classes to unique locations in the double vs. single space. Large dots denote the mean single and double frequencies for each class.
